# CRFalign: A Sequence-structure alignment of proteins based on a combination of HMM-HMM comparison and conditional random fields

**DOI:** 10.1101/2022.02.03.478675

**Authors:** Sung Jong Lee, Keehyoung Joo, Sangjin Sim, Juyong Lee, In-Ho Lee, Jooyoung Lee

## Abstract

We built a method of sequence-structure alignment (called CRFalign) which improves upon a base alignment model based on HMM-HMM comparison by employing pairwise conditional random fields (pCRF) in combination with nonlinear scoring functions of structural and sequence features. The total scoring function consists of a base scoring part based on HMM-HMM profile comparison plus additional nonlinear scoring part which is implemented by a set of gradient boosted regression trees. In addition to sequence profile features, various structural features are employed including secondary structures, solvent accessibilities, environment-dependent properties that give rise to position-dependent as well as environment-dependent match scores and gap penalties. Training is performed on reference alignments at superfamily levels or twilight zone chosen from the SABmark benchmark set. We found that our alignment method produce relative improvement in terms of average alignment accuracies, especially for the alignment of remote homologous proteins. We found that our alignment method produced (by using Modeller) better modeling results especially in the relatively hard targets compared with other methods. CRFalign was successfully applied to the stages of fold recognition and multiple sequence alignment in CASP11 and CASP12 competition on protein structure predictions.

## INTRODUCTION

Comparing a protein sequence with another sequence or a sequence with a known protein structure is one of the most important tasks in bioinformatics, especically in 3D structure modeling. In spite of striking new developments in recent years (such as Alphafold [1, 2]) on 3D protein structure modeling based on contact predictions via deep learning, sequence structure alignment method is still essential in protein fold recognition for finding templates for given target sequences. For example, in the case of Alphafold2, up to four templates are used as input to the structure modeling.

In traditional template-based modeling (TBM), the model qualities are highly dependent on finding the best templates and good alignments between the target sequence and the templates. When multiple templates are given, multiple alignments between the sequence and templates[3–6] are utilized. However, multiple alignment is strongly dependent on the alignment accuracies of pair-wise sequence-sequence or sequence-structure alignments. Also improving pairwise sequence-structure alignment is important for finding better templates for protein structure modeling.

Various kinds of profile comparison methods have been developed to improve the alignment quality between sequences. For example, there are several score functions available for calculating the match scores between profiles such as the dot product score [7], the Jensen-Shannon divergence score [8], the log average score [9], and the Pearson’s correlation score [10], etc. SparksX[11] builds on previous profile-profile (comparison) alignment methods of SP (SP1, SP2, SP3, SP4, SP5) series[12–15] by incorporating additional features incrementally, such as secondary structures and solvent accessibility with linear combination. HHpred on the other hand is based on comparison of HMM profiles[16], [17]. More recently, more discrimitive methods of conditional random fields have been applied to pairwise alignment and fold recognition. These include Contralign [18], BoostThreader[19, 20] and MRFalign [21]. BoostThreader, in particular, employs a nonlinear scoring function by means of regression trees, or neural networks. Among these methods, HHalign method of HHpred has been consistently successful with particularly fast performance.

In this work, we built a method for pairwise alignment between a sequence and a structure which combines an HMM-HMM comparison scoring scheme (HHalign, HHblits [22]) and an additional nonlinear scoring function based on pairwise conditional random fields with boosted regression trees. We incorporate boosted regression trees at each stage of the training steps with various features including profile-profile similarity, secondary structure similarity, similarity of solvent accessibility, as well as environmental features. These nonlinear scoring functions are expected to provide complicated relationships between neighboring features for propensities of match states or gaps.

We find that improvement of alignment accuracy can be achieved especially for pairwise alignments between protein sequence and structures with remote homologies. We evaluated the alignment quality of CRFalign by modeling the 3-D structures via Modeller for different test sets from SABmark benchmark databasei and some past CASP targets. Here, we found that the TM-score’s and RMSD scores of modelled structures from CRFalign showed consistently larger improvement over those from base model alignments for the case of the *hard* remote homolog targets.

## MATERIALS AND METHODS

Here we present the formalism of conditional random fields as applied to pairwise sequence-structure alignment of proteins [19]. For a given pair of protein sequences *s* (which we denote, in this work, as the sequence for the known *structure*) and *t* (which we denote as the sequence for the *target* with unknown structure), an arbitrary alignment between the structural sequence *s* and the target sequence *t* can be represented as a sequence of match (*M*), insertion (*I*) or deletion (*D*) states. If we suppose that *L_s_* is the sequence length of the *structure*, and that *L_t_* is the sequence length of the *target*, then this sequence of alignment states can also be represented as an alignment path on a rectangular lattice of dimensions *L_s_* × *L_t_* where a diagonal path corresponds to *M*, a horizontal one to an *I* and, a vertical one to a *D*. Here we assume that the structural sequence (*s*) lies along the horizontal with the sequence length *L_s_* and the target sequence (*t*) along the vertical with the sequence length *L_t_*.

Now let us denote the sequence of alignment states (with alignment length *L*) as *a* ≡ (*a*_0_, *a*_1_, *a*_2_, · · ·, *a_L_*, *a*_*L*+1_) where *a_i_* ∈ {*M, I, D*} with *i* = 1,· · ·, *L* in addition to *a*_0_ which indicates the **BEGIN** state as well as *a*_*L*+1_ denoting the **END** state. In the formalism of conditional random fields, a probabilistic model for the pairwise alignment is constructed where the probability *P*(*a*|*s, t*) of each of these alignments can be written as,

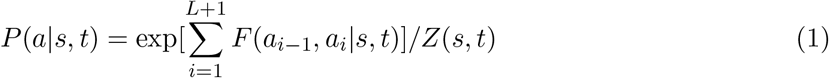

where the function *F*(*a*_*i*−1_, *a_i_*|*s, t*) represents the log-likelihood of the transition from the alignment state *a*_*i*−1_ at *i* − 1 to the next state *a_i_* at the alignment position *i* and *Z*(*s, t*) is the normalization factor with 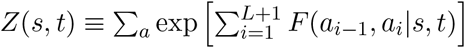 which is also called the partition function. The function *F*(*a*_*i*−1_, *a_i_*|*s, t*) corresponds to the alignment score at alignment position *i* in conventional pairwise sequence alignment. Here, however, it depends not just on the present alignment state *a_i_* but also on the previous alignment state *a*_*i*−1_ through the local structural or sequence features of residues located around the alignment position *i* − 1 and *i*. This makes it possible to naturally incorporate position dependence or environmental dependence in match scores as well as gap penalties. Important features include similarities of sequence profiles between the sequence and structure, similarities of secondary structures and solvent accessibilities. Originally, conditional random field formalism was based on the function *F* with linear combination of various scores **??**. J. Xu *et al* proposed a method based on nonlinear scoring functions for *F* such as neural networks or boosted regression trees [19, 20]. These nonlinear scoring functions can take nontrivial correlations between different features into account. Optimal choice of the functions *F* are obtained through training on a set of referece alignments such that the average probability of these reference alignments get maximal values.

In our alignment, *F* consists of a sum of successive scoring functions as follows,

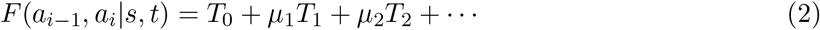

where *T*_0_ is a base alignment model of scoring function and *T_i_*’s (for *i* ≥ 1) are successive nonlinear scoring functions to be determined by optimization of the probabilities of occurrences of some reference alignment set. Suppose that *P*(*a*_*i*−1_, *a_i_*|*s, t*) refers to the net probability that the specific transition from the alignment state *a*_*i*−1_ at position *i* − 1 to the next state *a_i_* at position *i* occur (which is also called the posterior probability). Then it is straightforward to show that, for any alignment model defined by *F*(*a*_*i*−1_, *a_i_*|*s, t*)

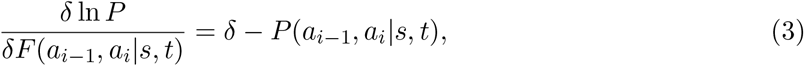

where *δ* is equal to 1 if *a*_*i*−1_ at position *i* − 1 and the next state *a_i_* at position *i* actually pass along a reference alignment and if not, it is equal to zero. A simple way to understand this relation is that, when the alignment model defined by *F*(*a*_*i*−1_, *a_i_*|*s, t*) is optimal (i.e., at maximum), then the right hand side of Eq. (3) should be zero, in other words, *P* (*a*_*i*−1_, *a_i_*|*s, t*) should be equal to 1 (maximal probability) for the states on the alignment path (*δ* = 1), and equal to zero for the states not on the alignment path (*δ* = 0). Here *P*(*a*_*i*−1_, *a_i_*|*s, t*) is obtained by summing over all the alignments of *s* and *t* with the restriction that a specific transition *a*_*i*−1_ to *a_i_* should occur at specific position for the pair of sequences. This can be easily computed by Forward and Backward algorithm [23]. Now, we can see that the successive scoring functions *T_k_*, *k* = 1, 2, · · · can be constructed by any machine learning methods for the functional gradient of the ln *P* (for the alignment in question to occur) with respect to *F*, which can be easily sampled from training alignments using the right-hand-side of Eq. (3). Here in this work, we implemented *T_k_* as boosted regression trees, which is fast and efficient.

In previous works by Xu, et al [19], the beginning alignment model *T*_0_ for *F* was chosen as the trivial alignment model with *F*_0_(*a*_*i*−1_, *a_i_*) = 0 for all possible transitions at all positions, which roughly corresponds to a random alignment model where all possible pairs of residues have equal probability of alignment as well as equal probabilities for all gaps. Here in our work, instead of beginning with the random alignment model, we chose to begin with some reasonable alignment method which is already available and build our full alignment model by adding nonlinear scorning functions within the framework of conditional random fields. Here, we chose a scoring scheme adapted from HHalign[16] as the base scoring scheme.

HHalign is a pairwise alignment method based on comparison of HMM profiles of protein sequences[16]. In order to construct a pairwise comparison of two HMM’s, HHalign introduces five pair-alignment states which are *MM*, *MI*, *IM*, *GD*, *DG* where *M* denotes a Match state in the HMM of a specific residue position for the structure sequence or the target sequence, *I* an Insertion state, *D* a deletion state, and finally *G* denoting a Gap state. For a given alignment between two HMM’s, the alignment score of HHalign consists of HMM-HMM profile match score, transition probability score and secondary structure score as follows.

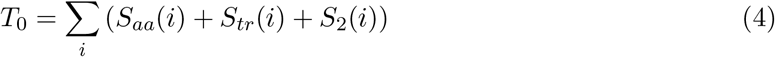

where *S_aa_*(*i*) denotes the similarity score between the HMM profiles of the two columns at the alignment position *i*, *S_tr_* represents the propensity of allowed transitions at the alignment position *i* that are transitions between a pair state and itself and between pair state *MM* and pair states *MI*, *IM*, *DG* or *GD*. The last term *S*_2_(*i*) represents the similarity score between the secondary structure of the template (structure) residue and the predicted secondary structure of the target residue. Usually, for the template structure, the secondary structure information from DSSP[24] is used, while, for the target residue, secondary structure prediction from PSIPRED [25].

Accommodation of HHalign-type scoring into our CRF alignment model with additional non-linear scoring function may be implemented in two different ways that are called in this work as three-state scheme or five-state scheme. In the three-state scheme of CRF alignment model, we reduce the five states *MM*. *MI*, *IM*, *DG*, *GD* of HHalign to the usual three states *M*, *I*, *D* via the reduction mapping of (*MM* → Match(*M*), *MI* → Insertion(*I*), *IM* → Deletion(*D*), *DG* → Insertion(*I*), *GD* → Deletion(*D*)). Note that *MI* and *DG* are both reduced to the same alignment state of *I*, while *IM* and *GD* to *D*. For a given alignment path in this three-state scheme, the zeroth order scoring is obtained by reduction of the HHalign scores such that for an Insertion (*I*) or a Deletion (*D*) state, the larger score of the two corresponding states in the original five-state model from HHalign is chosen.

It is also possible to construct a CRF alignment model with five-state scheme (i.e., without reduction to the three states) incorporating the full HHalign scoring and additional nonlinear scoring functions. And, in this scheme of five state, for the purpose of training from the reference alignments, it is necessary to assign appropriate five-state label for each of the alignment position between the target and the template. Since the reference alignments are built by some structure alignments without relation to the HMM profiles of the targets or templates, it is not straightforward to assign five state labels to the alignment states due to the ambiguity between *DG* vs. *MI* as well as *GD* vs. *IM*. One possible solution to this problem is to choose the unique assignment along the reference alignment path for which the HHalign score becomes maximum. We implemented and tested both the three-state model and the five-state model.

Once the base alignment model for three-state is fixed, additional nonlinear scoring functions (*T*_1_, *T*_2_, · · ·) can be constructed via training on a set of reference alignments as follows. One can evaluate the right-hand side of the Eq. (3) based on the base alignment model *T*_0_ to get the functional derivative 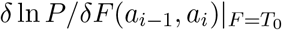 for any pairwise alignment. Now, we sample transitions from the set of reference training alignments. Both positive samples (i.e., those transitions appearing in the training alignments) as well as negative samples (i.e., those transitions that are not appearing in the training alignments) are taken. For these samples, one can compute the right-hand side of the Eq. (3) 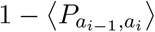. These target values together with the relevant input features can now be used to train the first additional scoring function *T*_1_ i.e., first correction to the alignment model, where any machine learning methods can be employed. The constant factor *μ*_1_ can be chosen to control the degree of greediness of the training. For training these gradients, we used so-called gradient boosted regression trees [26]. Also, the partition function *Z*(*s, t*) can be calculated using the standard Forward-Backward algorithm for the given alignment model. Now, when this training is completed for *T*_1_, we are now equipped with a first order corrected alignment model, which can again be used for training the next order regression trees *T*_2_ for further correction, using new samples evaluated at *T*_0_ + *μ*_1_ · *T*_1_. The constants *μ_i_* (*i*=1, 2, 3, · · ·) are weight parameters that can be adjusted to control the degree of convergence in the training, where we chose *μ_i_* = 0.2 for all *i* in this work.

We consider the following input features for the regression trees of the function *F*.

A. Features for match states: We assume that the *i*-th position of the template (structure) sequence is aligned (i.e., matched) to the *j*-th position of the target sequence.
  1. Similarity between the HMM-profile of the structure and that of the template, which is calculated as the logarithm with base 2 of the dot product between the HMM-profiles of *s* and *t*. The HMM profiles are obtained from “Hblits” developed by Söding, et al.

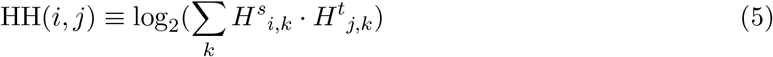

where *H^s^_i,k_* denotes the *k*-th component of the hmm profile (position specific scoring matrix) at the residue position *i* of the structure template, and *H^t^_j,k_* denotes the *k*-th component of the hmm profile at the residue position *j* of the target.
  2. Secondary structure similarity : We employ three-state classification for secondary structures, i.e. *C* (=Coil), *H* (=*α*-Helix), and *E* (=*β*-strand). For the template structures *s*, each residue position is assigned by DSSP [24] one of these three states. For the targets, we use the 3-state secondary structure prediction of PSIPRED which gives three values of relative likelihood of secondary structures corresponding to (*C*, *H*, *E*). For a match between the observed secondary structure in 3-state representation at *i*-th position of the template structure and the predicted secondary structure at *j*-th position of the target, the secondary structure similarity is determined as the component value of the predicted secondary structure at *j*-the position of the target corresponding to the observed secondary structure at *i*-th position of the template structure as follows

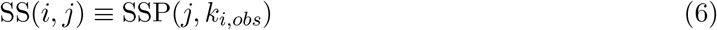

where *k_i,obs_* denotes the observed secondary structure (one of *C*, *H*, or *E*) at the residue poition *i* of the template, and SSP(*i, k_i,obs_*) denotes the corresponding predicted (PSIPRED) value of the secondary structure at the residue position *j* of the target.
  3. Solvent accessibility : As for the solvent accessibility feature, we use a three-state classification scheme with labels of ‘Buried’, ‘Intermediate’, and ‘Exposed’ states. These are determined by the values of the relative solvent accessibility (RSA) of a specific residue with ranges 0%-9% (’Buried’), 9%-36% (’Intermediate’), and 36%-100% (’Exposed’). For the template structure, the solvent accessibility state is determined by DSSP. Also for the case of target (query) sequence, we employ the program SANN [27] which predicts RSA propensity with 3-state scheme for a sequence with unknown 3D structure. For a match between the observed RSA (obtained from DSSP) at *i*-th position of the template and the predicted RSA (from SANN) at *j*-th position of the target, the feature value is chosen as

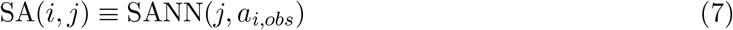

where *a_i,obs_* denotes the observed RSA (one of the three states ‘Exposed’, ‘Buried’, or ‘Intermediate’) at the residue poition *i* of the template (from DSSP), and SANN(*j, a_i,obs_*) denotes the corresponding predicted value of the RSA at the residue position *j* of the target.
  4. BLOSUM matrix, Gonnet matrix, and Kihara matrix : For a match state between *s* at position *i* and *t* at position *j*, we take as additional features the corresponding matrix elements of the BLOSUM62 matrix [28] *B*(*s_i_, t_j_*), Gonnet250 matrix [29] *G*(*s_i_, t_j_*), and the Kihara matrix [30] *K*(*s_i_, t_j_*).
  5. Environmental fitness score : For the residue position *i* of the template, DSSP provides the secondary structure and the solvent accessibility for the residue. The environmental fitness score for the match between *i*-th residue position of the template and *j*-th residue position of target is obtained from the weighted average of the environmental fitness potential of the template’s local environment state with the position specific frequency matrix (PSFM) *H^t^*(*j, k*) derived from the HMM profile of the target as follows

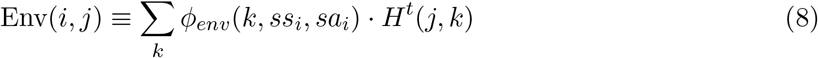

where *ϕ_env_*(*k, ss_i_, sa_i_*) denotes the environmental fitness potential of amino acid *k* for the secondary structure state *ss_i_* and solvent accessibility state *sa_i_* of the template at *i*-th residue position (which was borrowed from the PROSPECT II [31]).
  6. Neighborhood similarity score : For the match between *i*-th position of the template and *j*-th position of the target, this score measures the similarity between the neighboring residues within a fixed window. Suppose we set the window size *n_w_* ≡ 2 ∗ *f* + 1 with *f* ≥ 1, then the neighborhood similarity between the template and the target at offset position *k* from (*i*,*j*) is defined as the sum of Pearson correlations of the PSFM, SS and SA at (*i* + *k*)-th residue position of the template and the (*j* + *k*)-th residue position of the target with *k* = 0, ±1, ±2, · · ·, ±*f*.

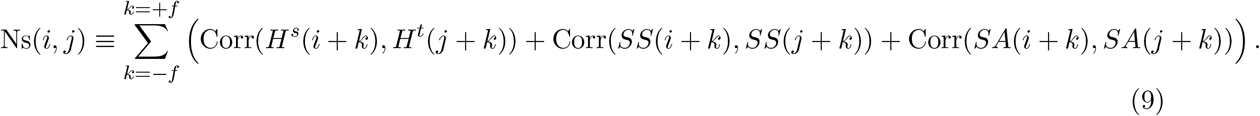 Here Corr(*x, y*) denotes the Pearson correlation of vectors *x* and *y* with the same length of components which is defined as

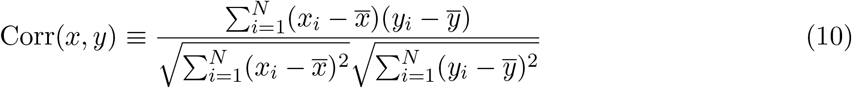

where 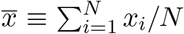 is the average value of *x_i_*, *i* = 1,. . . , *N*. As for the size of the neighboring window, we set *n_w_* = 9 (i.e. *f* = 4).
B. Input features for the Gap states:
  1. Seven-component reduced-profile features derived from the HMM profile based on seven classes of residues: For a gap state (both at the template and target), the corresponding inserted residue was classified into seven classes as (i) class *I* of hydrophobic and aliphatic residues including Ala, Ile, Leu, and Val. (ii) class *II* of hydrophobic and aromatic residues including Phe, Trp, and Tyr. (iii) class *III* of polar residues including Asn, Cys, Gln, Met, Ser, and Thr. (iv) class *IV* of Acidic charged residues including Asp, and Glu. (v) class *V* of Basic charged residues including Arg, His and Lys. (vi) class *V I* of the residue Gly. (vii) class *V II* of the residue Pro. Suppose there is a gap at position *i* in the template with the corresponding residue at position *j* of the target, one can construct a simple reduced profile by collecting and summing those component values of the PSFM (derived from the HMM profile) of the target residue at *j*-th position, such that those components belonging to the same class are summed over. That is,

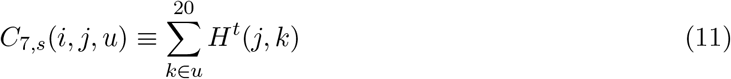

where *H^t^*(*j, k*) denotes the *k*-th component of the HMM profile (position specific scoring matrix) at the residue position *i* of the target. The seven-class index *u* ranges from the class *I* to the class *V II* as indicated above. Similar method can be applied to the case of a gap in the target.
  2. Secondary structure : For a gap state at the target (i.e., insertion at the template), the observed secondary structure (from DSSP) of the template at the corresponding inserted residue is used as the secondary structure feature, which is represented in a 3-vector form with the Coil (*C*) state corresponding to (1, 0, 0), the Helix (*H*) state to (0, 1, 0) and the Extended Beta (*E*) to (0, 0, 1). On the other hand, for a gap state at the template (i.e., insertion at the target), the predicted secondary structure propensity with three components (from PSIPRED) of the target at the corresponding inserted residue is used as the secondary structure feature.
  3. Solvent accessibility : Similarly to the case of secondary structure, for a gap state at the target (i.e., insertion at the template), the observed solvent accessibility in three states (derived from DSSP) of the template at the corresponding inserted residue is used as the input feature. The three states of “Buried” (*B*), “Intermediate” (*I*), and “Exposed” (*E*) are represented as (1, 0, 0), (0, 1, 0), and (0, 0, 1) respectively. On the other hand, for a gap state at the template (i.e., insertion at the target), the predicted solvent accessibility in three states (from SANN) of the target at the corresponding inserted residue is used as the input feature.
  4. Local gap propensity from secondary structure environment: For a gap state at the target (i.e., insertion at the template), the predicted secondary structure information (from PSIPRED) of the target in the seven neighboring residues (i.e. from −3 to +3 separation from the gap position) is combined to give the Coil (or Loop) propensity as

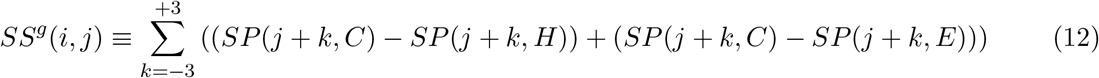

where *SP* (*j* + *k, C*) denotes the component of the predicted secondary structure at the residue position *j* + *k* of the target, to be found in a Coil state *C*, *SP* (*j* + *k, H*) the component of the predicted secondary structure to be found in a Helix state *H*, and similarly *SP* (*j* + *k, E*) the component for the Beta strand state *E*. On the other hand, for a gap state at the template (i.e., insertion at the target), we use similar formula on the neighboring residues for the template except that here, instead of the predicted secondary structure, the observed secondary structure is used, which is represented in a 3-vector form with the Coil (*C*) state corresponding to (1, 0, 0), the Helix (*H*) state to (0, 1, 0) and the Extended Beta (*E*) to (0, 0, 1).
  5. Local gap propensity from solvent accessibility environment: For a gap state at the target (i.e., insertion at the template), the predicted solvent accessibility information (from SANN) of the target in the seven neighboring residues (i.e. from −3 to +3 separation from the gap position) is combined to give the Coil (or Loop) propensity as

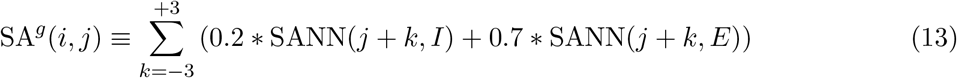

where SANN(*j* + *k, I*) denotes the component of the predicted solvent accessibility at the residue position *j*+*k* of the target, to be found in an “Intermediate” state *I*, SANN(*j*+*k, E*) the component of the solvent accessibility to be found in an “Exposed” state *E*. Hence we put more weight on the exposed residues than buried or intermediate residues. Similarly for a gap state at the template (i.e., insertion at the target), we use similar formula on the neighboring residues for the template except that here, instead of the predicted solvent accessibility, the observed solvent accessibility (in normalized 3-vector form) is used. Therefore, the summand in the above formula becomes 0.2 for the state “Intermediate” and 0.7 for the “Exposed” state.
  6. Additional features for indicators of the end position of the sequences: In addition to the above input features for the gap states, we also use separate indicators for the end positions of each of the two sequences for special treatment of end gaps. That is, we define

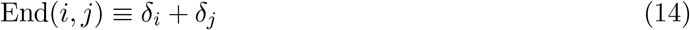

where *δ_i_* (*δ_j_*) = 1 if *i* (*j*) is at the beginning (N-terminal) or at the end (C-terminal) of the sequence and 0 otherwise. In the CRF alignment model, two kinds of alignments are possible. One is the so-called Viterbi alignment algorithm which selects the one highest scoring alignment (i.e., highest probability). The other method of alignment is the so-called MAP (MAximum Posterior Probability) alignment which first calculates the net probability *P*(*s_i_, t_j_*) for a specific pair of residues *s_i_* (of the template structure) and *t_j_* (of the target) may align in all possible alignments, and then find, through a standard dynamic programming, the alignment that optimizes the sum of these values without gap costs,

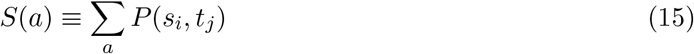

where the sum is performed over all pair matches. This is also called as Maximum Accuracy Alignment (MAC). MAP alignment tends to produce more true matches compared with the Viterbi alignment (see the section on Results).

In contrast to global alignment (where the alignment begins at the first residues of the target and the template), local alignments can be generated by allowing the alignment to begin (and end) at any position of the target and the template without scoring for the end gaps. In the case of MAP alignment, this can be conveniently implemented by introducing a threshold value *m_th_* for the match which can range from 0 to 1 and subtract *m_th_* from *P* (*s_i_, t_j_*) for all matches as follows,

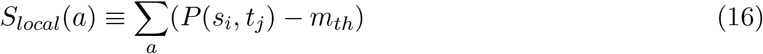

where, in the dynamic programming, the alignment can begin at any position of the target and template with no costs for end gaps. As for internal gaps, additional penalties of −0.5 ∗ *m_th_* are added to avoid unnatural internal gaps being produced. A special case of alignment mode which is called *glocal* (*glocal+local*) alignment is commonly adopted, where the alignment can begin (and end) at internal positions of either the target or the template but not *both*. If we suppose that the sequence of the structure template runs along horizontal axis on top boundary of a rectangular lattice, with that of the target running along vertical axiz on the left boundary, this corresponds (in terms of the alignment path) to beginning the alignment on the upper or left boundaries of the rectangle and ending on the opposite sides (bottom or right boundaries) in the dynamic programming.

## RESULTS AND DISCUSSION

As for the reference alignment set for training our sequence structure alignment method, we chose the SABmark (version 1.65) benchmark set [32] which was designed to assess protein sequence alignment algorithms, especially for the case of remote homologous pairs of proteins. SABmark consists of two sets of pairwise and multiple alignment sets with high resolution X-ray structures derived from the SCOP classification. These sets, Twilight Zone and Superfamilies, are known to cover the entire known fold space with sequences very low to low, and low to intermediate similarity respectively.

The Twilight Zone set consists of 209 sequence groups that each represent a SCOP fold. Sequence similarity is very low with the sequence identities ranging between 0-25% and also with the structures being distantly similar. SABmark homepage states that “This set therefore represents the worst case scenario for sequence alignment, which unfortunately is also the most frequent one, as most related sequences share less than 25% identity” [32]. On the other hand, the Superfamilies set consists of 425 groups, each of which representing a SCOP superfamily. The sequence pairs share at most 50% identity. Even though this set in general consists of less difficult pairs (than the Twilight Zone) they still represent challenging problems for sequence alignments.

We chose three sets of reference alignment each consisting of 200 pairwise alignments from the Superfamilies set and from the Twilight Zone. These three sets are labelled as NG200, NF200, and TW200 respectively. These are chosen in such a way that the pairs are evenly distributed among different groups of families, so that as many groups of families as possible are covered. Among these three sets, two of them (NG200 and NF200 set) are from the Superfamilies set with average sequence identities of 24.2%, 21.2%, respectively. The remaining set of TW200 is derived from the Twilight zone set with average sequence identity of 13.8%. By training our sequence structure alignment methods on these sets with different levels of sequence similarity, we may be able to compare the modeling capabilities of the resulting alignment methods and choose the most efficient one among those results.

HMM files were generated by using hhmake tools of HMM hhsuite [16] [22]. In this work, for the case of three state scheme, the nonlinear scoring functions *T*_1_, *T*_2_, · · · are implemented by the so called boosted regression trees. At each step of training, e.g., *T_i_* with *i* = 1, 2, · · ·, there are three different boosted regression trees, one for each of the three states at the current position: match (*M*), insertion (*I*) and deletion (*D*) state, respectively.

For each pairwise alignment of the reference training set, we take each of the alignment states along the reference alignment path as a positive sample (for training the scoring function). If it is a match state (*M*), we add the set of corresponding features together with the target label value 1 − *P*(*a*_*i*−1_, *a_i_*) in Eq. (3) to the sample set for the boosted regression tree for the match state. Similarly for insertion (*I*) or deletion (*D*) states along the reference alignment path, we add the corresponding features and the label value to the boosted regression trees for *I* and *D* respectively. Now, we have to also collect negative samples, that is, those transitions that do not appear on the reference alignments. Suppose that the sequence length of the structure *s* and the target *t* are *L_s_* and *L_t_* respectively. And if we let the alignment length to be *L_a_* then we have *L_a_* ≤ *L_s_* + *L_t_*. Then we can see that (e.g., for three state alignment model) there are about ≃ 3(*L_s_* · *L_t_* − *L_a_*) negative samples which is usually much larger than the alignment length *L_a_*. We randomly selected some integer (*N_f_*) times the alignment length *L_a_* for the size of the negative samples, where we took *N_f_* = 16. That is, we took 16 · *L_a_* transitions that are not on the alignment path. These transitions were distributed evenly among the three alignment states *M*, *I*, and *D*. We tried other values for *N_f_*, but the present value was found to be most effective in terms of the training accuracy and training time. This resulted in around 2 · 10^5^ samples for each of the three states. At each step of the training, the boosted regression trees consist of six regression trees with each tree having a depth of five. As for the choice of the parameters *μ_k_*, as mentioned above, we simply set *μ_k_* = 0.2 for all steps *k*. Change of this parameter did not show much difference in the performance of the resulting alignment model.

For each of the above three sets (NG200, NF200, TW200), in order to perform training of our alignment model and then perform validation test in terms of alignment accuracies, we divided the set into four subsets of 50 pairs each, and then carried out a four-fold training and test with 150 pairs for training and the remaining 50 pairs for test in turn.

Figure 1 shows the training and test accuracies (with Viterbi scoring) for these three sets as the training step increases. We see that on all three cases, the training and the test accuracies on average improve up to a certain regression steps but after about five to seven regression steps (depending on the sets), the accuracies begin to decrease, even though there are cases when the test alignment alignment accuracies continue to improve further up to the last step (step 9). Figure 2 shows the relative improvements of the CRFalign test accuracies over the zeroth order model for the three sets. We can clearly see that, for the hard alignment set of NF200 and TW200 as compared with the easier set of NG200, more improvement is achieved especially in terms of the Viterbi alignment accuracies.

**FIG. 1:**
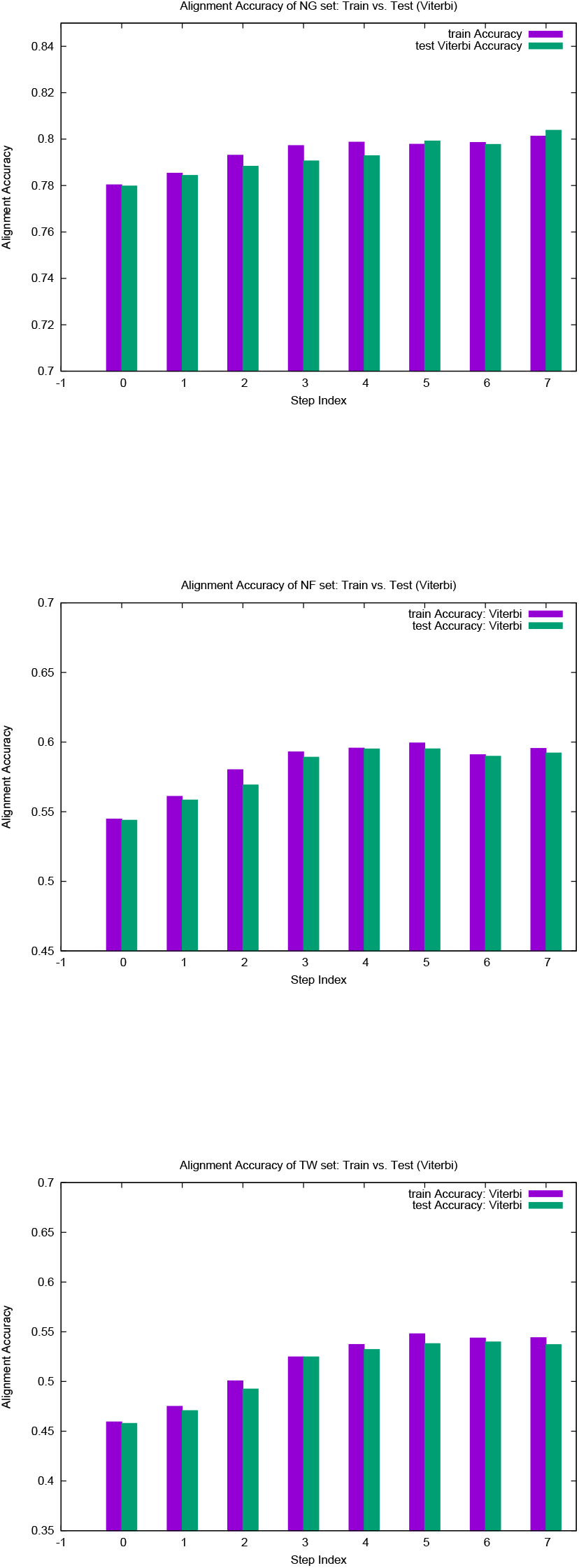
Training and test accuracies for the Viterbi alignment on the three sets (a) NG, (b) NF, and (c) TW of 200 reference alignments from SABmark.

**FIG. 2:**
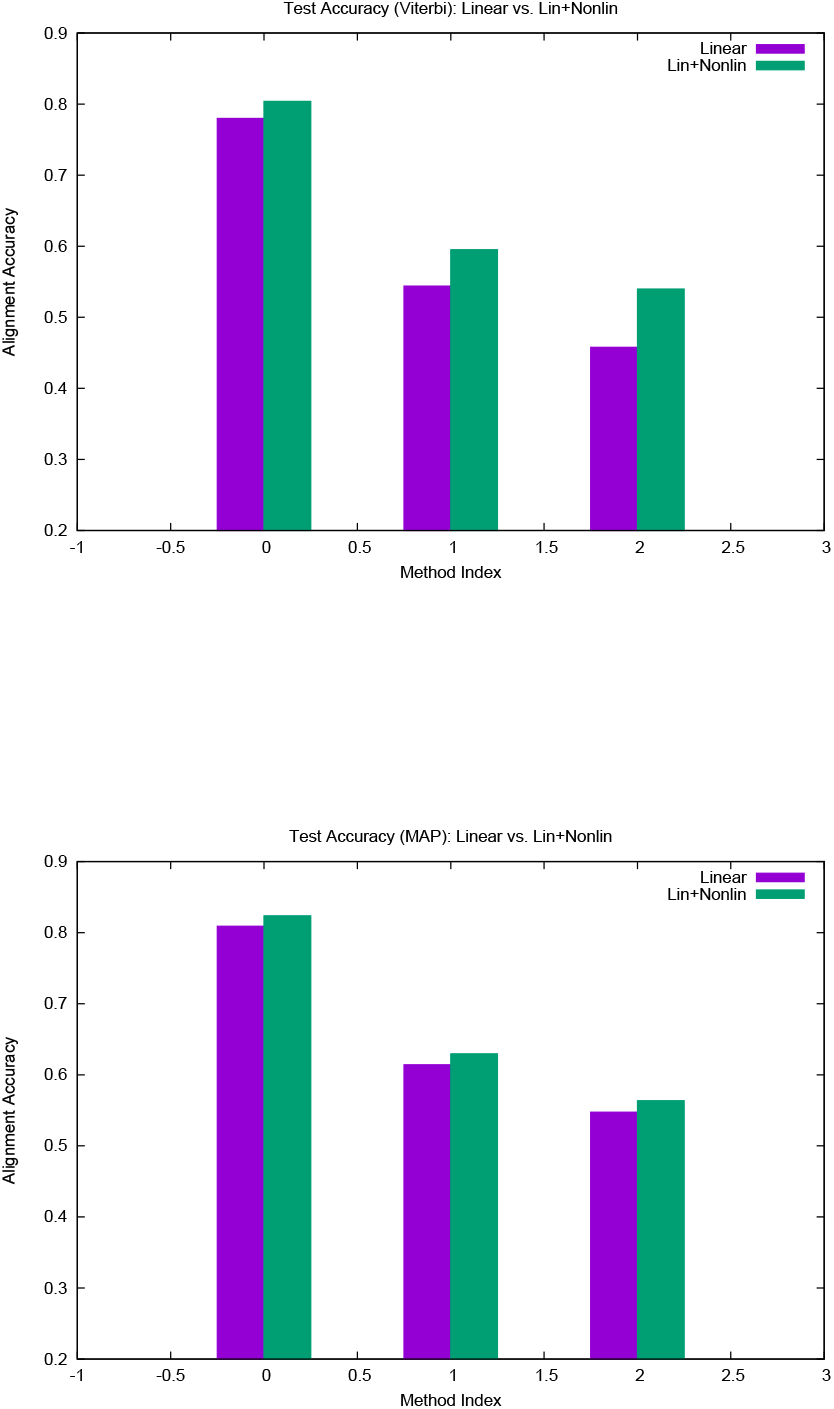
Training and test accuracies at Maximum for (a) the Viterbi alignment as well as (b) the MAP alignment on the three sets (NG, NF, and TW) of 200 reference alignments.

In order to assess our alignment method in terms of protein structure modeling for proteins in the SABmark set, we chose the whole 200 pairs of the NF200 set to train our alignment model and then performed sequence structure alignment with the trained model together with structure modeling on two independent test sets using Modeller program based on the alignment. One of the two test sets called NG64 consists of 64 pairs chosen from the Superfamilies set of the SABmark. On the other hand, the second test sets called TW55 consists of 55 pairs chosen from the Twilight Zone set of the SABmark, representing more difficult alignment situations. Both set consist of pairs of sequences that are less than 30 % sequence identity against those of the training set NF200 with the average of the maximum sequence identity against the training set being 16.6% (NG64) and 16.3% (TW55) respectively.

Figure 3 shows the modeling results on NG64 and TW55 test sets comparing CRFalign method with HHalign, where we see that notable improvements were made by CRFalign over HHalign results. Table I shows the average TM-score for the modeling results where again we find that for the hard set of TW55 the improvement was bigger. Average TM score of NG64 set by CRFalign was 71.96% while that for HHalign was 71.39%. On the other hand, the average TM score of TW55 set by CRFalign was 48.83% while that for HHalign was 46.32%.

**FIG. 3:**
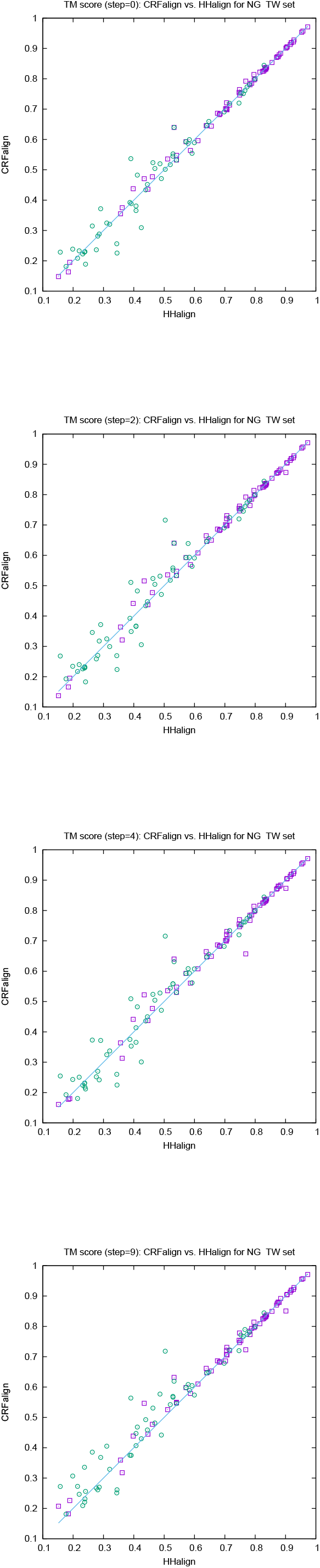
TM-scores of structure models of NG set and TW set obtained by running Modeller on the CRFalign alignments at various train steps (steps 0, 2, 4, and 9) in comparison with the results for the HHalign.

**TABLE I:** Modeling accuracies in TM-score of the test sets NG64 and TW55 based on training with NF200

| Set | HHalign | Base model | CRFalign |
| --- | --- | --- | --- |
| NG64 | 0.7139 | 0.7176 | 0.7196 |
| TW55 | 0.4632 | 0.4669 | 0.4883 |

Figure 4 shows one example where CRFalign result was fed into Modeller with the resulting model exhibiting significant improvement over that of HHalign. Shown is the 3D structure of the chain A of d1nr0a1 (which is the seven-bladed beta propeller domain of C. elegans actin-interacting protein 1) with the template d1fwxa2 (d1nr0a1-d1fwxa2, TM*_ref_* = 75.38%, ID = 9.4%). Note that the sequence ID to the template sequence is only 9.4% but still CRFalign in combination with Modeller could produce a structure of TM-score with TM ≃ 71.8% which is close to the optimal TM-score limit of ≃ 75.38%. In contrast hhalign could not properly close the propeller shaped domain with relatively poor value of the TM-score of TM*_hha_* = 50.36%. Figure 5 shows the alignment between the template and our target sequence from which we can see that there are a few large gaps in the alignment which would be difficult to correctly align without the help of structure based features and nonlinear scoring model for the alignment. Another example is shown in Fig. 6 where *β* protein (The Outer Membrane Protein OMPX from E. Coli 1qj8a) is illustrated based on the alignment (d1qj8a-d1i78a TM*_ref_* = 72.65%, ID = 3.4%) which could roughly produce the correct fold pattern with TM*_CRF_* = 56.38% as compared with hhalign which failed in producing the correct *β* patterns on one side with TM*_hha_* = 39.02. In this case the sequence identity is even lower with ID = 3.4%. The alignment to the template is shown in Fig. 7 where we again find regions of big gaps. We can compare with the optimal structure (referece) alignment. Final example is shown in Fig. 8 which shows the structure of d1hnja2 (Beta-Ketoacyl-acyl carrier protein synthase III) with the alignment d1hnja2-d1hnja1 (TM*_ref_* = 59.88%, ID=10.4 %). Here, CRFalign could produce TM*_CRF_* = 57.68% which is quite close to the ideal value of TM*_ref_* = 59.88%. In contrast HHalign could produce the model with TM*_hha_* = 48.62% only, failing to reproduce much of the secondary structural elements. Figure 9 shows the alignment by CRFalign.

**FIG. 4:**
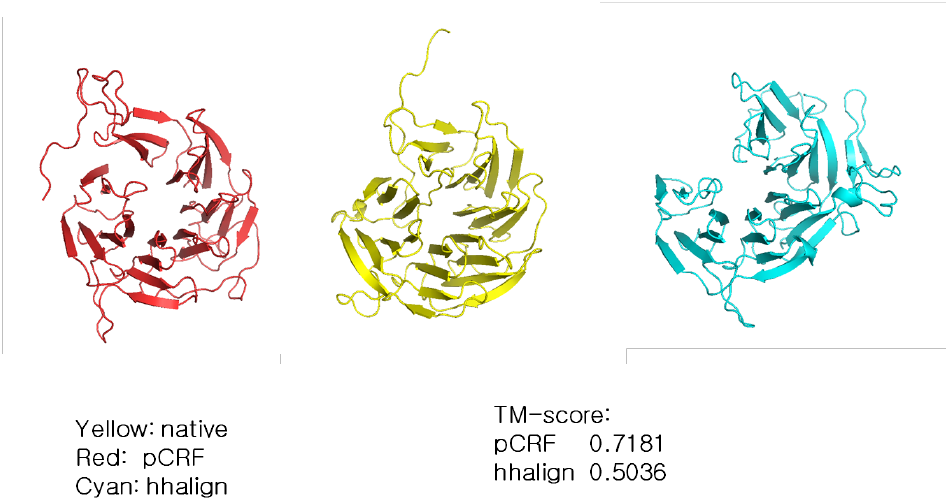
d1nr0a1-d1fwxa2 (*TM_ref_* = 0.75381, *ID* = 9.4%) C.elegans actin-interacting protein1 Seven-bladed beta-propeller domain, Yellow: native Red: CRFalign Cyan: hhalign, TM-score: *CRF* − > 0.7181, *hhalign*− > 0.5036.

**FIG. 5:**
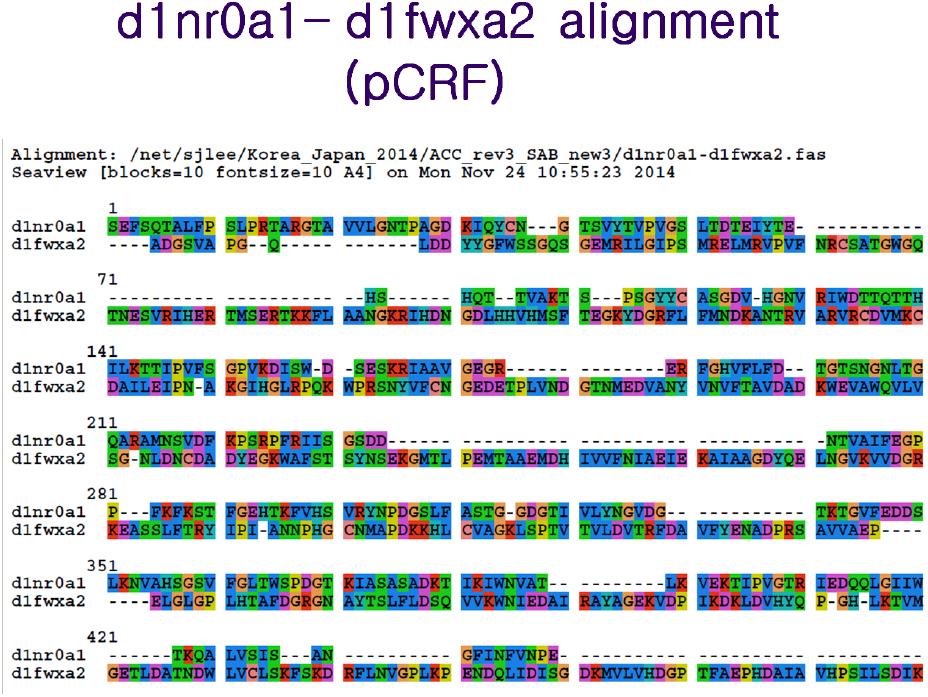
d1nr0a1-d1fwxa2 alignment, CRFalign alignment.

**FIG. 6:**
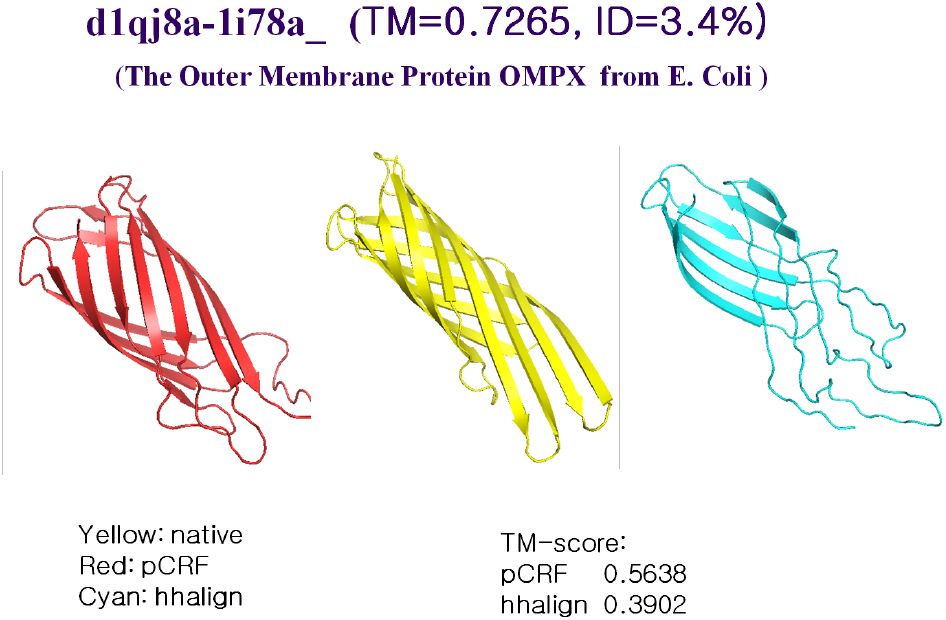
d1qj8a-d1i78a (TM=0.7265, ID=3.4 %) The Outer Membrane Protein OMPX from E. Coli. Yellow: native Red: CRFalign, Cyan: hhalign, TM-score: *CRF* = 0.5638, *hhalign* = 0.3902

**FIG. 7:**
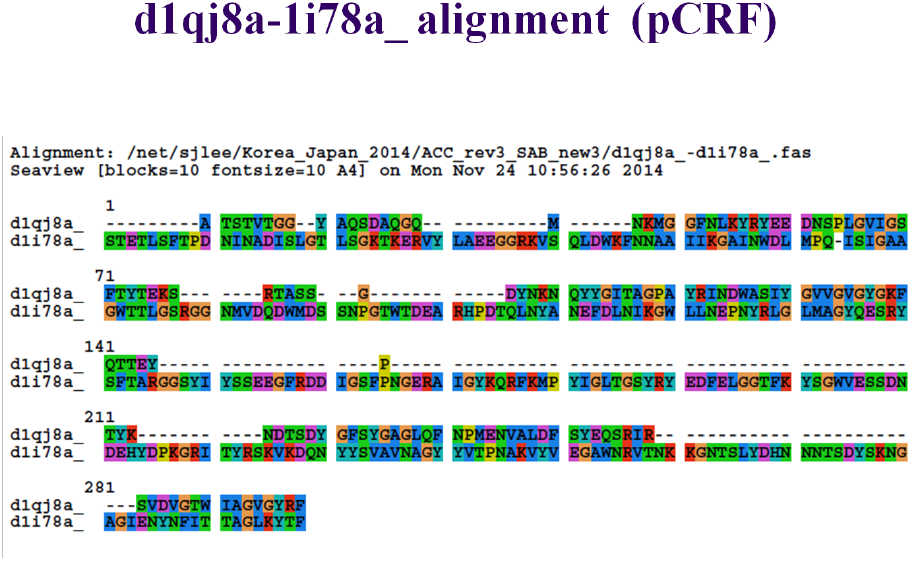
d1qj8a-d1i78a alignment, CRFalign alignment

**FIG. 8:**
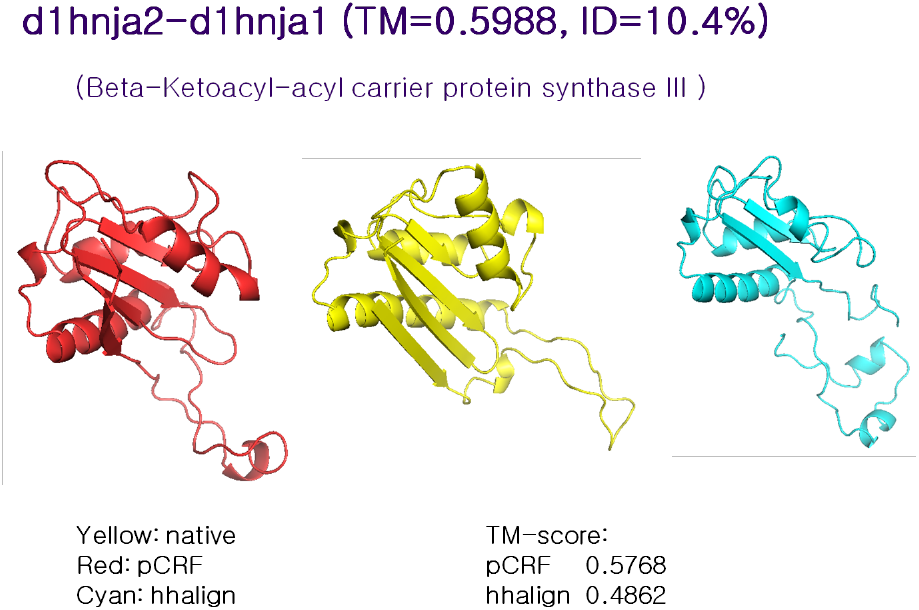
d1hnja2-d1hnja1 (*TM_ref_* = 0.5988, *ID* = 10.4%) Beta-Ketoacyl-acyl carrier protein synthase III, Yellow: native Red: CRFalign Cyan: hhalign, TM-score: *CRF* = 0.5768, *hhalign* = 0.4862

**FIG. 9:**
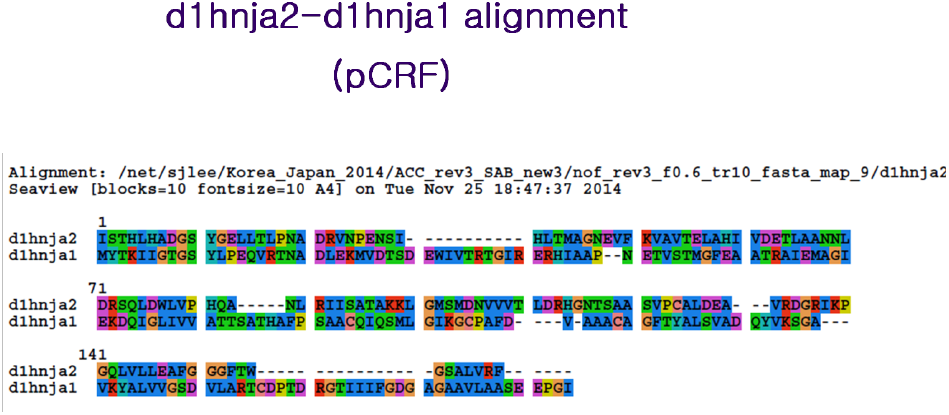
d1hnja2-d1hnja1 alignment, CRFalign

For building working alignment models (targeted for blind structure prediction such as CASP), we trained several hundred different alignment models (i.e., accumulting different sets of boosted regression trees) on the three sets (NG200, NF200, and TW200) using the whole 200 pairwise alignments for each of the three sets. In order to choose optimal alignment models, we need to test these for their modeling capabilities. For this, we tested these on CASP10 targets with appropriate templates by performing alignment and modeling.

We chose 58 single-domain targets from CASP10, for which there exist templates. Among these, for 50 of them, we could choose two templates. Hence in all, we have 108 pairs to align and model. One of the optimal models was chosen from training on NG200 set with three state alignment scheme and the training step of 4 which we call BRT-NG4. We also obtained other optimal alignment models with comparable performance with five-state alignment scheme, but we chose the three-state scheme BRT-NG4 mainly because three-state scheme is about twice faster than the five-state alignment.

Figure 10a shows the comparison of the TM-scores for the modeled structures with the base alignment model vs. HHalign where we see that the average TM-score with the base alignment model (TM*_base_* ≃ 0.5246) is lower than that of the HHalign model (TM*_HHalign_* ≃ 0.5286). However, on the right side, Fig. 10b shows the comparison between the CRFalign (with BRT-NG4) result and HHalign, where we see various targets for which the base model gave relatively poor result are now showing appreciable improvement with the resulting average TM-score of TM*_CRFalign_* ≃ 0.5321.

**FIG. 10:**
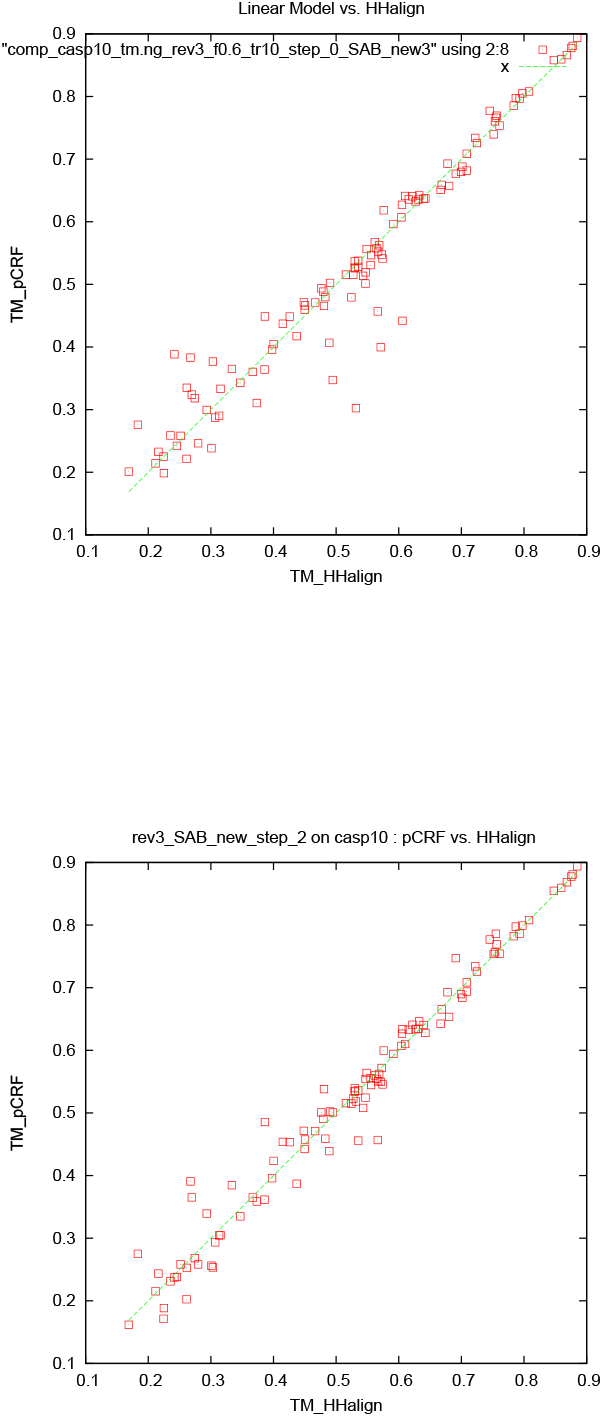
Comparison of TM-scores for CASP10 targets by Modeller modeling based on (a) base alignment (average *TM* − *score* = 0.5246) vs. HHalign alignment (average *TM* − *score* = 0.5286) and (b) full CRF alignment (average *TM* − *score* = 0.5321) vs. HHalign.

We also tested the five-state model with a new training set called TR367. The training set TR367 consists of 367 pairs of proteins from SABmark superfamily (299 pairs) and twilight zone set (68 pairs). In order to assess the pair-wise alignments of five-state alignment model, we prepared two test sets W200 and S200 from SABmark benchmark set. W200 set consists of 200 pairs chosen from the twilight zone subset of SABmark set, while those of S200 are 200 pairs from the superfamily set, where all the sequences in the test set have sequence identities less than 20% against those sequences in the training set (TR367). Therefore, pairs in W200 set should be considered, in general, harder (i.e., remote homologs) than those of S200 set. Figure 11a-c shows the training and test Accuracy of alignments for TR367 training set and W200 test set as well as S200 test set. We can see here also that the average alignment accuracy in the W200 set shows larger relative improvement due to pCRF than that in the case of S200 set (table II).

**FIG. 11:**
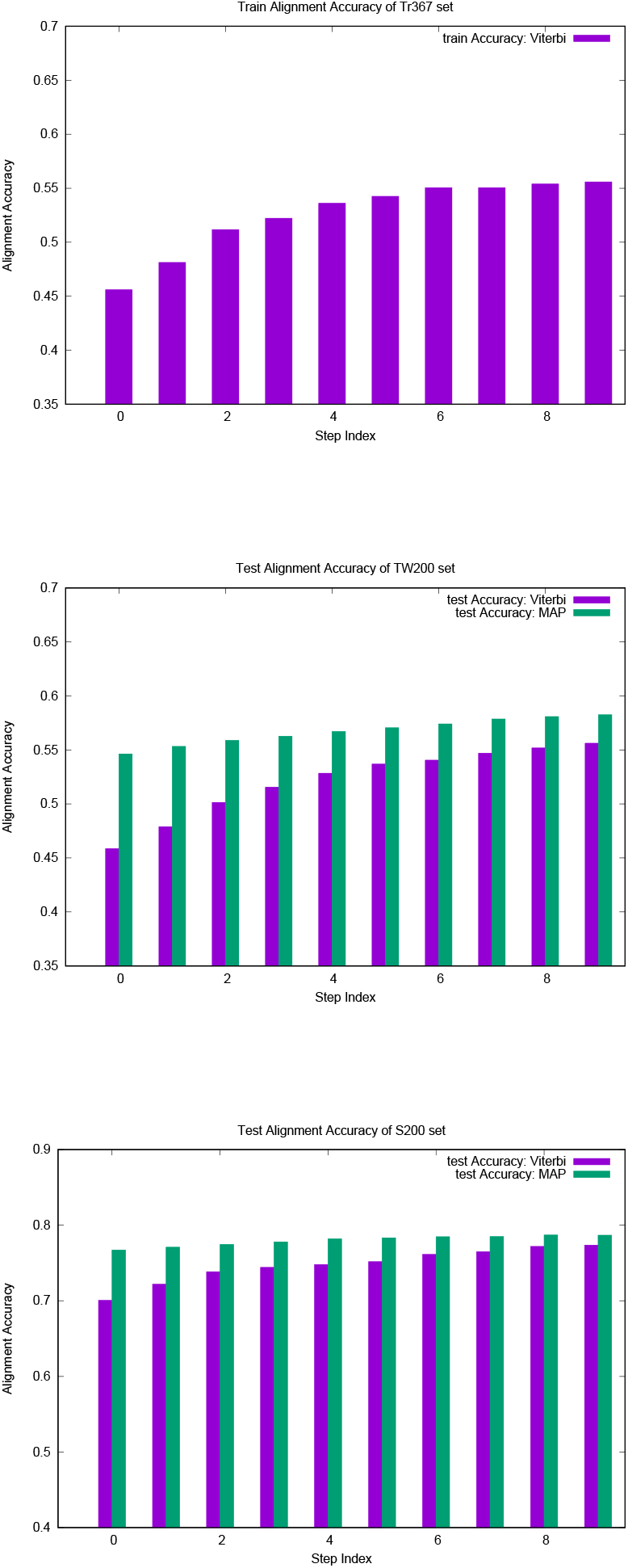
(a) Training alignment accuracy on TR367 set with five-state model training (above) (b) Test alignment accuracies on W200 set with the same five-state model. Both viterbi alignment and MAP alignment accuracies are shown (middle) (c) Test alignment accuracies on S200 set with the same five-state model (below).

**TABLE II:** Alignment accuracies in the test sets W200 and S200 based on training with TR367

| Set | Base model | CRFalign |
| --- | --- | --- |
| W200 | 0.546 | 0.583 |
| S200 | 0.767 | 0.787 |

Table 3 shows the modeling accuracies on the two test sets W200 and S200. We can recognize larger improvement in the TM score in the case of W200 set compared with that of S200 set which is consistent with the alignment accuracies shown above.

**TABLE III:** Modeling accuracies in TM-score of the test sets W200 and S200 based on five-state training with TR367 set

| Step number | W200 | S200 |
| --- | --- | --- |
| 0 | 0.4329 | 0.5792 |
| 1 | 0.4350 | 0.5812 |
| 2 | 0.4366 | 0.5808 |
| 3 | 0.4396 | 0.5809 |
| 4 | 0.4416 | 0.5826 |
| 5 | 0.4431 | 0.5827 |
| 6 | 0.4463 | 0.5820 |
| 7 | 0.4470 | 0.5833 |
| 8 | 0.4494 | 0.5849 |
| 9 | 0.4495 | 0.5870 |

Figures 12a-b through Fig. 16a-b show examples of the significantly improved model structures from the W200 set using five-state models with pCRF as compared with those from the base model.

**FIG. 12:**
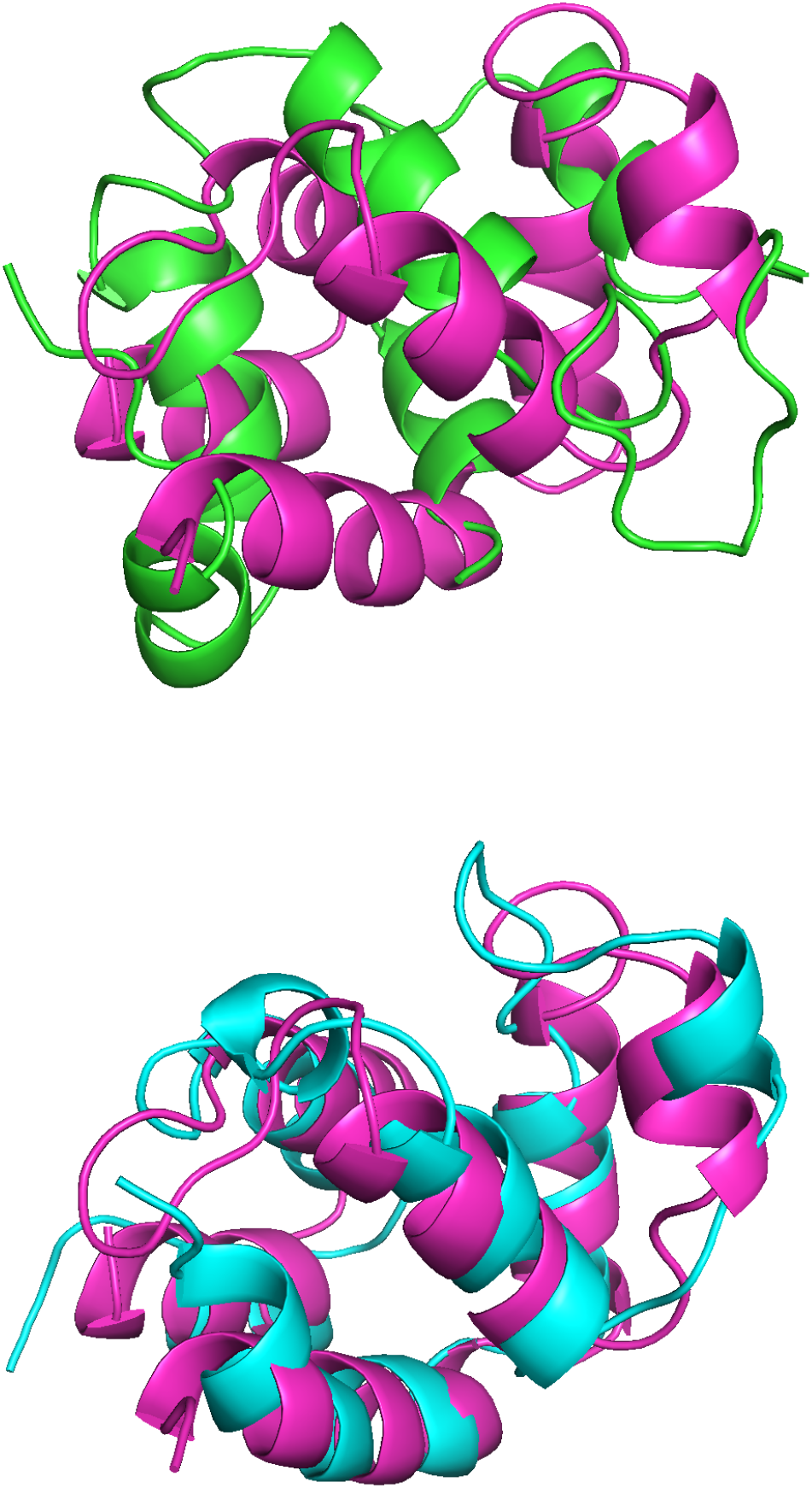
(a) Structure model based on d1a1w-d1dgna base alignment (green) overlapped with the experimental structure of d1a1w (red purple) (with *rmsd* = 11.97 *Å*) and (b) that based on CRFalign alignment (cyan) overlapped with the experimental structure (red purple) (with *rmsd* = 2.156*Å*).

**FIG. 13:**
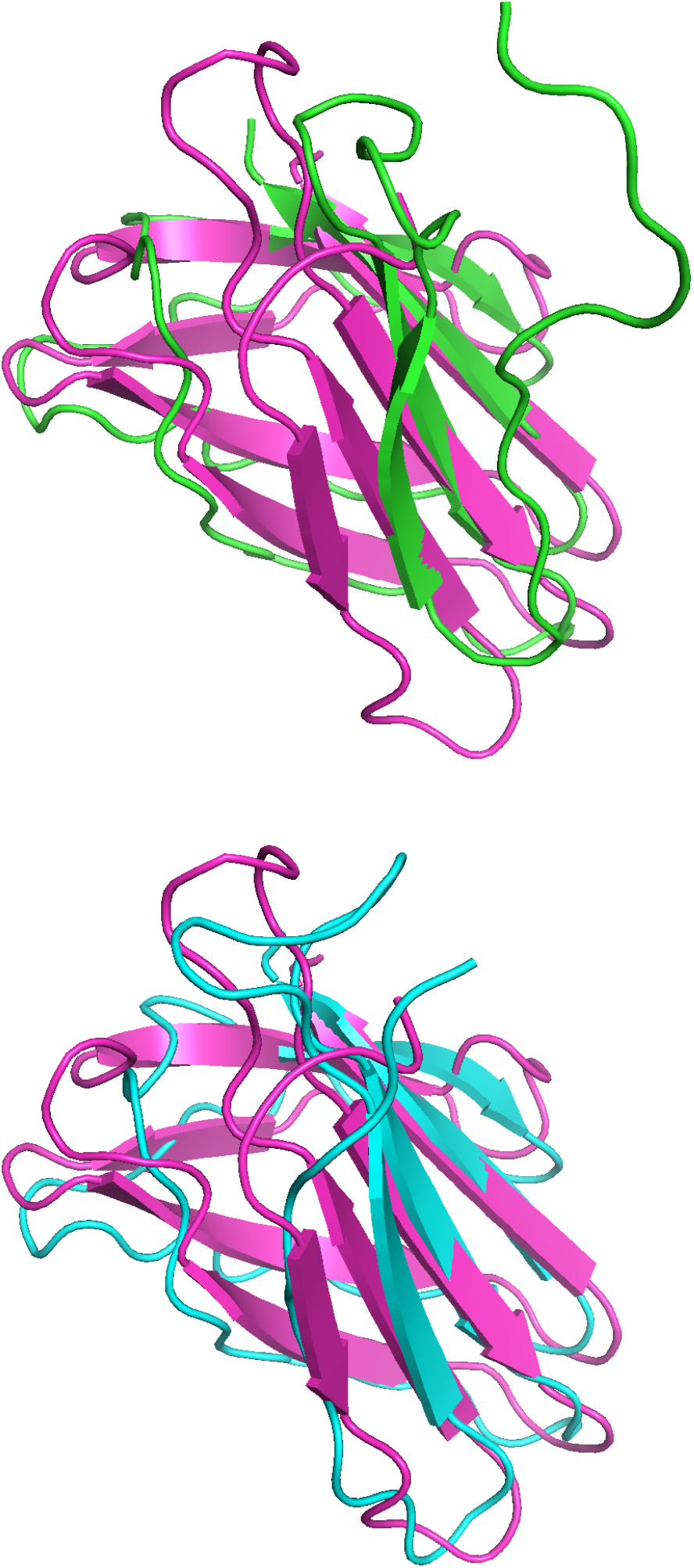
(a) Structure model based on d1e43a1-d1m7xa2 base alignment (green) overlapped with the experimental structure of d1e43a1 (red purple) (with *rmsd* = 3.486*Å*) and (b) that based on CRFalign alignment (cyan) overlapped with the experimental structure (red purple) (with *rmsd* = 2.378*Å*).

**FIG. 14:**
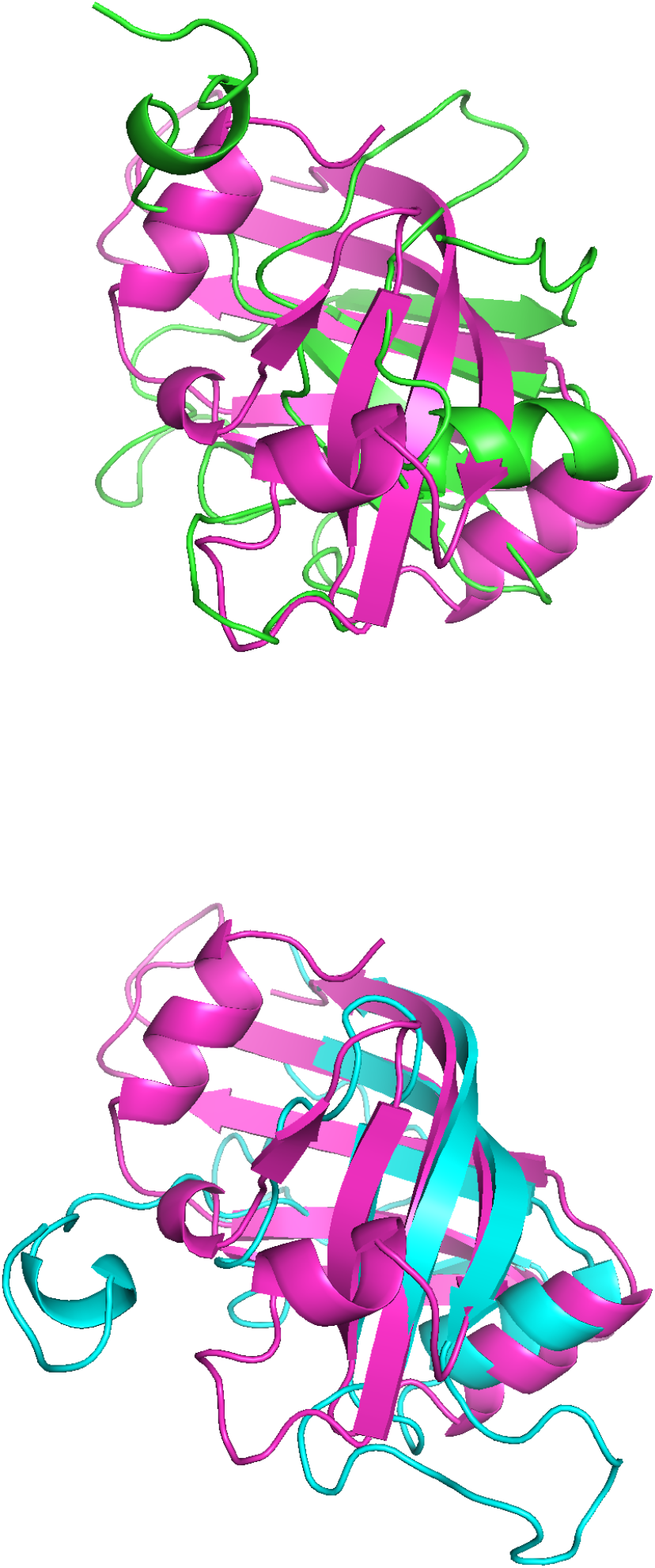
(a) Structure model based on d1flma-d1ejea base alignment (green) overlapped with the experimental structure of d1flma (red purple) (with *rmsd* = 15.223*Å*) and (b) that based on CRFalign alignment (cyan) overlapped with the experimental structure (red purple) (with *rmsd* = 2.387*Å*).

**FIG. 15:**
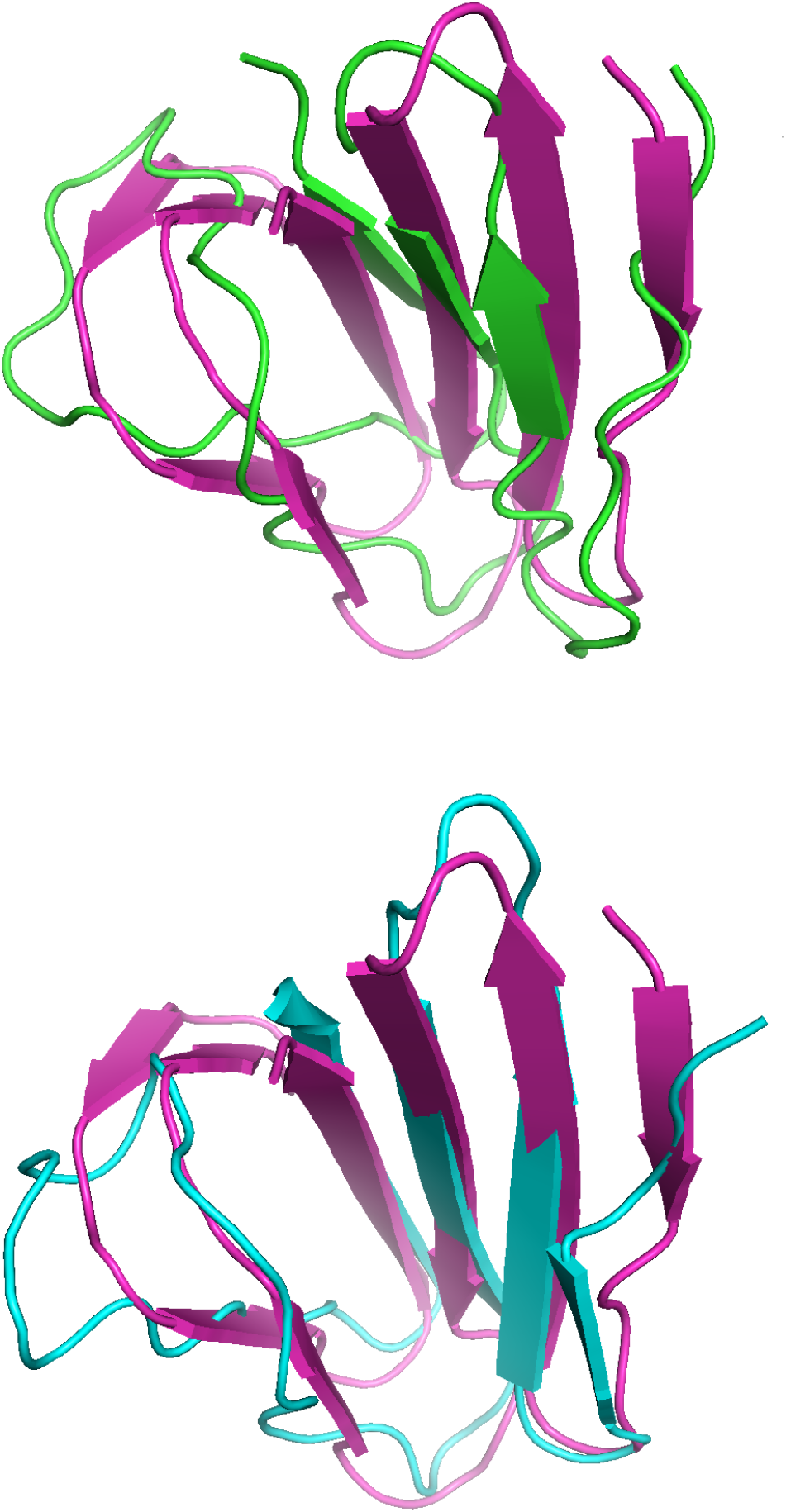
(a) Structure model based on d1gjwa1-d1ktba1 base alignment (green) overlapped with the experimental structure of d1gjwa1 (red purple) (with *rmsd* = 4.616*Å*) and (b) that based on CRFalign alignment (cyan) overlapped with the experimental structure (red purple) (with *rmsd* = 2.348*Å*).

**FIG. 16:**
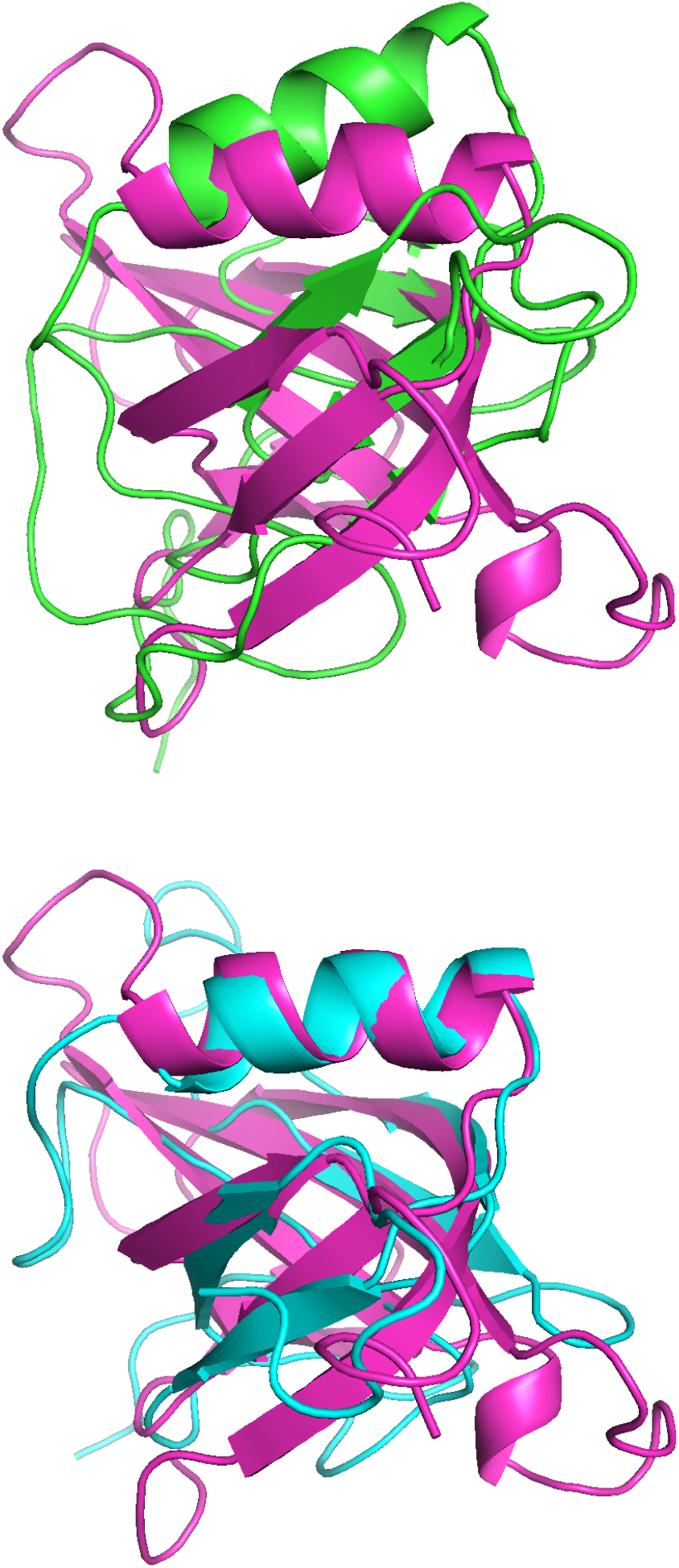
(a) Structure model based on d1prtd_-d1prtb1 base alignment (green) overlapped with the experimental structure of d1prtd (red purple) (with *rmsd* = 14.425*Å*) and (b) that based on CRFalign alignment (cyan) overlapped with the experimental structure (red purple) (with *rmsd* = 2.540*Å*).

## CONCLUSIONS

A sequence-structure alignment method CRFalign is presented that improves upon a reduced three-state or five-state scheme of HMM-HMM profile alignment model by means of conditional random fields with nonlinear scoring on sequence and structural features implemented with boosted regression trees. CRFalign can extract and exploit complex nonlinear relationships among sequence profiles and structural features including secondary structures, solvent accessibilities, environment-dependent properties that give rise to position-dependent as well as environment-dependent match scores and gap penalties. Training of the CRFalign is performed on a chosen set of reference paiwise alignments from the SABmark benchmark set which consists of Twilight Zone set and Superfamilies set with pairs of sequences very low to low, and low to intermediate sequence similarity respectively. We found that our alignment method produce significant improvement in terms of average alignment accuracies, especially for the alignment of remote homologous proteins. Comparison of the modeling capabilities of our alignment on independent pairs of SABmark set with those of HHalign showed that our alignment method produced better modeling results especially in the relatively hard targets. CRFalign was successfully applied to the initial stages of fold recognition and as an input to the multiple sequence alignment called (MSACSA) in the recent CASP11 and CASP12 competition on protein structure predictions.

## ACKNOWLEDGEMENTS

This research was supported by Basic Science Research Program through the National Research Foundation of Korea (NRF) funded by the Ministry of Science and ICT (NRF-2017R1E1A1A01077717 and NRF-2018R1D1A1B07049312). We thank KIAS Center for Advanced Computation for providing computing resources.

